# Reconstructing intra-tumor fitness landscapes from scSeq CNA genotypes via simulation-based Bayesian inference and Deep Learning

**DOI:** 10.64898/2026.04.10.717795

**Authors:** Maryam KafiKang, Pavel Skums

## Abstract

Inferring the selective effects of copy-number alterations (CNAs) from clonal tumor data is essential for understanding tumor evolution. In practice, intra-tumor evolutionary parameters are typically estimated by fitting population genetic models to observed data using maximum likelihood or Bayesian methods. However, realistic mechanistic models often lead to intractable likelihoods, limiting the applicability of conventional inference approaches.

Here, we introduce a likelihood-free, simulation-based framework for inferring intra-tumor selection coefficients directly from clonal CNA profiles. Our approach employs neural posterior estimation to amortize inference across simulated tumors and uses normalizing flows to flexibly parameterize high-dimensional posterior distributions while enabling robust uncertainty quantification.

Our primary model, CloneMLP-NPE, learns representations of whole-tumor CNA genotypes using a multilayer perceptron (MLP)-based encoder. We compare this model against two baselines: (i) a Set Transformer encoder applied to the same whole-tumor CNA profiles, and (ii) a consensus-based approach that relies only on the CNA profile of the most abundant clone. On held-out simulations, CloneMLP-NPE achieves the strongest overall performance, yielding well-calibrated posterior distributions and more accurate posterior mean estimates than both baselines.

## 1 Introduction

Cancer progression is widely understood as an evolutionary process. Neoplasms often arise from a single founding cell, and continued genetic instability generates heritable variation on which selection can act, producing successive expansions of fitter subclones and increasing intratumor heterogeneity [6]. In many cancers, this evolutionary “search” is driven not only by point mutations but also by large-scale copy-number alterations (CNAs), including whole-chromosome or chromosome-arm gains and losses, which reshape dosage-sensitive pathways and recur across tumor types [5].

Within this framework, selective pressure is manifested through differential clonal growth. Clones carrying advantageous CNA configurations expand relative to others, whereas clones with deleterious configurations contract. Quantifying these selective effects is computationally challenging, yet it is essential for understanding the mechanisms underlying cancer progression [15].

The problem becomes more tractable when intra-tumor populations can be observed at multiple time points [9]. In practice, however, most available datasets consist of single snapshots rather than longitudinal measurements. Over the past decade, a number of methods have been proposed to infer cancer fitness landscapes from bulk and single-cell sequencing data [4, 10, 13]; nevertheless, the problem remains far from solved.

Inference of intra-tumor selection is most commonly approached through phylogenetic or phylodynamic models [10, 11]. While these methods make the problem more computationally tractable, they typically rely on simplifying assumptions about the evolutionary process. More recently, there has been increasing interest in parameter inference using more realistic, simulation-based tumor evolution models, together with likelihood-free and Bayesian calibration techniques [5, 16]. Similar strategies have also become increasingly important in other scientific domains, particularly in physics [1].

These mechanistic simulators generally induce likelihoods that are difficult or impossible to evaluate directly. A common workaround is to use likelihood-free strategies such as Approximate Bayesian Computation (ABC), which compares simulated and observed data through summary statistics rather than explicit likelihood evaluation. Recent work has applied ABC to infer chromosome- and arm-level selection parameters under chromosomal instability (CIN), demonstrating both the promise of simulator-based inference and the practical challenges imposed by summary-statistic design and computational scaling as the parameter space grows [5, 16]. In parallel, modern simulation-based inference (SBI) has advanced substantially through the use of learned conditional density estimators that approximate posteriors from simulated training pairs, enabling amortized inference and principled uncertainty quantification even in settings with intractable likelihoods [1, 12].

In this work, we present a likelihood-free Bayesian framework for inferring chromosome arm-level selection coefficients directly from clonal CNA profiles. We adopt neural posterior estimation (NPE), a conditional density-estimation approach to SBI, and we parameterize the posterior with expressive normalizing flows [7, 8, 12]. Our primary model, which we refer to as **CloneMLP-NPE**, uses the whole-tumor CNA matrix containing CNA profiles for all subclones in a tumor and learns a representation of this matrix using a multilayer perceptron (MLP)-based encoder prior to posterior estimation. To assess the value of this representation, we compare it against two alternative baselines: **CloneAtt-NPE**, which applies a Set Transformer encoder to the same whole-tumor CNA matrix, and **DominantClone-NPE**, which uses only the CNA profile of the largest clone. By combining a mechanistic simulator with amortized posterior learning from clonal CNA data, our framework aims to recover posterior distributions over CNA selection parameters while preserving, to varying degrees, information about tumor clonal composition. Finally, we evaluate all models on held-out simulations using posterior diagnostic analyses to assess both inference quality and uncertainty calibration.

## 2 Methods

### 2.1 Simulator: SISTEM

We used SISTEM (SImulation of Single-cell Tumor Evolution and Metastasis) [14], a Python framework for simulating tumor growth, metastasis, and DNA sequencing data under genotype-driven selection. Unlike neutral coalescent models or simple birth–death simulators, SISTEM models somatic clonal selection within an agent-based framework and can generate mutation profiles, read counts, and synthetic sequencing reads together with ground-truth lineage and migration information.

SISTEM simulates cancer cell populations across one or more anatomical sites over discrete generations, where each cell is characterized by its anatomical location and a genome representation that includes allele-specific copy-number profiles and single-nucleotide variants (SNVs) [14]. At the end of a cell’s lifespan, the cell either dies or replicates into two daughter cells, which may acquire additional mutations; surviving cells may also migrate between anatomical sites. Replication probability is determined by cellular fitness, which depends on both genotype and anatomical site and is scaled according to a predefined population growth model. Selection is represented through a multiplicative fitness landscape specific to each anatomical site [14].

In this study, we used SISTEM’s copy-number-profile (CNP) selection formulation, in which cellular fitness is computed directly from copy-number states [14]. In particular, SISTEM provides a chromosome-arm selection model that assigns a selection coefficient to each chromosome arm, with each coefficient reflecting the relative balance of oncogenes and tumor suppressor genes on that arm. To generate evolving CNA landscapes over time, SISTEM simulates SNVs together with multiple CNA mechanisms, including segmental gains and losses, chromosome-arm and whole-chromosome missegregation, and whole-genome duplication events.

### 2.2 Simulation procedure and parameter sampling

We generated training pairs by simulating clonal tumor evolution with SISTEM under its chromosome-arm selection model [14]. For each sampled simulator setting, the full parameter vector *ϕ* comprised 46 parameters: two CNA event-rate parameters (`arm_rate and chromosomal_rate`) and 44 chromosome-arm selection coefficients for the autosomal arms. In the inference task, we aimed to recover only the 44 chromosome-arm selection coefficients, denoted by *θ* ∈ ℝ^44^. The two CNA event-rate parameters were therefore treated as nuisance simulator inputs rather than inference targets.

For each sampled parameter setting, we generated 25 independent simulation replicates to capture stochastic variability in the evolutionary process. Each replicate produced a clonal population from which we extracted chromosome-arm CNA summaries to construct the observation *x* (Section 2.3).

In total, we sampled 2,500 simulator parameter settings and generated 25 independent replicates for each, resulting in 62,500 simulated tumors overall. The dataset was split at the level of parameter settings into training and test sets containing 80% and 20% of the sampled settings, respectively.

#### Parameter sampling

For each simulation, we sampled the chromosome-arm CNA rate `arm_rate` and the whole-chromosomal CNA rate `chromosomal_rate`, which in SISTEM govern the per-generation probabilities of acquiring a chromosome-arm CNA and a whole-chromosome CNA, respectively [14]. Specifically, we sampled `arm_rate` ∼Unif(10^−5^, 10^−4^) and `chromosomal_rate` Unif∼ (10^−6^, 10^−5^).

We also sampled chromosome-arm selection coefficients { *δ*_*a*_} for the autosomal arms *a* ∈ { 1*p*, 1*q*, …, 22*p*, 22*q*} according to *δ*_*a*_𝒩∼ (0, 0.2), and supplied these values to the chromosome-arm selection model through an arm-indexed coefficient map.

#### Simulation settings

We fixed several SISTEM parameters across all simulations. We set `region_len` to 5 × 10^6^, which determines the size (in base pairs) of the uniform genomic regions used in SISTEM’s discretized genome representation [14]. We set `focal_driver_rate` to 5 × 10^−4^, corresponding to the per-generation probability of acquiring a driver focal (segmental) CNA [14]. We set `max_distinct_driv_ratio` to 0.9, a viability-checkpoint parameter, and set the minimum detectable population size `min_detectable` to 5 × 10^6^ cells, so that simulation at a given anatomical site terminated once the population reached at least `min_detectable` cells [14]. All remaining parameters were kept at their SISTEM default values [14].

#### CNA summaries

For each completed simulation, we summarized chromosome-arm copy-number alterations using SISTEM’s arm-level region counts. For a given clone, the arm-level copy-number ratio was defined as the total number of region copies on that arm divided by the number of discretized regions assigned to the arm. These chromosome-arm ratios were flattened in a fixed autosomal order (chr1–chr22) and normalized as normalize(*x*) = log_2_(max(*x*, 10^−3^)) − 1.

For each simulated tumor, we extracted two CNA-based representations. First, we derived a dominant-clone CNA profile, defined as a 44-dimensional normalized arm-level copy-number vector from the most populous clone. This representation was used for **DominantClone-NPE**. Second, we constructed a whole-tumor CNA matrix by collecting the normalized 44-dimensional chromosome-arm profiles of all clones from the first anatomical site into an *N*× 44 matrix, where *N* denotes the number of clones. This whole-tumor representation was used as input to both **CloneMLP-NPE** and **CloneAtt-NPE**. In this way, the dominant-clone representation provides a simplified single-clone summary, whereas the whole-tumor matrix retains information about intratumoral clonal heterogeneity.

### 2.3 Data representation

Using the chromosome-arm CNA summaries described above, we constructed tumor-level representations from the simulated clonal populations. Because a simulated tumor can contain a very large number of cells, we first grouped cells sharing the same normalized chromosome-arm profile into clones and recorded the frequency of each clone within the tumor population.

The clones were then ranked by frequency, and the 100 most frequent clones were retained for each tumor to obtain a compact representation. For each retained clone, we appended its relative frequency as an additional feature. As a result, each clone was represented by a 45-dimensional vector consisting of 44 chromosome-arm CNA features and 1 frequency feature.

This procedure produced a whole-tumor CNA matrix

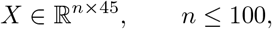

where each row *x*_*i*_ ∈ℝ^45^ corresponds to one retained clone. This whole-tumor matrix was used as input to both **CloneMLP-NPE** and **CloneAtt-NPE**.

In addition, for **DominantClone-NPE**, we extracted only the 45-dimensional feature vector corresponding to the most frequent clone. Thus, while the whole-tumor representation preserves information from multiple clones and their relative abundances, the dominant-clone representation provides a simplified summary based solely on the largest clone.

### 2.4 Simulation-based posterior inference

Each training example consists of a simulated tumor representation *X* together with the chromosome-arm selection coefficients *θ*∈ℝ^44^ used to generate it. Our goal is to learn an amortized approximation to the posterior

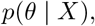

where *θ* denotes the 44 chromosome-arm selection coefficients and *X* is the observed tumor CNA representation.

We perform likelihood-free Bayesian inference using neural posterior estimation (NPE) with a normalizing-flow density estimator [7, 8, 12]. This approach enables direct approximation of the conditional posterior from simulated training pairs without requiring tractable likelihood evaluation. In contrast to point-estimation methods, NPE provides full posterior distributions over the selection coefficients, allowing us to quantify uncertainty in the inferred parameters.

To reduce variability arising from stochastic tumor evolution, we generated *T* = 25 independent simulation replicates for each sampled parameter vector *θ* and treated them as repeated observations from the same underlying parameter setting. Each replicate was first mapped to a fixed-dimensional embedding, and the embeddings of the 25 replicates associated with the same parameter vector were then aggregated by mean pooling to produce a single context vector *h*(*X*). This context vector was used to condition the normalizing-flow posterior model, which was trained on paired samples 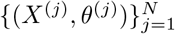.

Thus, all models in our study, namely **CloneMLP-NPE, CloneAtt-NPE**, and **DominantClone-NPE**, share the same inference pipeline: a tumor representation is encoded into a fixed-dimensional context vector, replicate-level contexts are aggregated by mean pooling, and the resulting representation is used to condition a normalizing flow that approximates the posterior over chromosome-arm selection coefficients.

### 2.5 Tumor encoders and baseline models

The models considered in this study differ only in how the simulated tumor representation is encoded before posterior estimation. For clarity, we refer to the proposed MLP-based model as **CloneMLP-NPE**, the Set Transformer baseline as **CloneAtt-NPE**, and the dominant-clone baseline as **DominantClone-NPE**.

#### CloneMLP-NPE

Our primary model uses the whole-tumor CNA matrix as input, where each row corresponds to one retained clone and includes 44 chromosome-arm CNA features together with the clone’s relative frequency. This matrix is encoded using an MLP-based encoder to produce a fixed-dimensional embedding for each simulation replicate. The resulting embedding is then used as the context vector for posterior inference.

#### CloneAtt-NPE

As a first baseline, we replaced the MLP encoder with a Set Transformer [3] applied to the same whole-tumor CNA matrix. This model preserves the same multi-clone input representation as **CloneMLP-NPE**, but uses attention-based set encoding to model interactions among clones while respecting permutation invariance.

#### DominantClone-NPE

As a second baseline, we used only the CNA profile of the most frequent clone in each simulated tumor. This dominant-clone representation is a single 45-dimensional vector containing 44 chromosome-arm CNA features and one clone-frequency feature. It provides a simplified alternative to the whole-tumor representation by ignoring the remaining clonal composition and intratumoral heterogeneity.

In all three cases, the encoder output is mapped to a fixed-dimensional context representation and passed to the same normalizing-flow posterior model. This design isolates the effect of the tumor representation and encoder architecture while keeping the downstream Bayesian inference procedure unchanged.

### 2.6 Posterior evaluation

We evaluated the learned posterior *q*(*θ*|*X*) on held-out simulated tumors for all three models—**CloneMLP-NPE, CloneAtt-NPE**, and **DominantClone-NPE**—using three quantitative criteria: posterior mean recovery, Z-score calibration, and posterior contraction.

#### Posterior mean recovery

We compared the posterior mean *µ*_*i*_(*X*) = 𝔼 [*θ*_*i*_ |*X*] to the ground-truth value 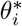 for each of the 44 chromosome arms. To quantify point-estimate recovery across held-out simulations, we report both the coefficient of determination *R*^2^ and the Pearson correlation coefficient between posterior means and true values. These two metrics provide complementary information: *R*^2^ evaluates how accurately posterior means recover the true parameter values relative to a mean baseline, capturing both bias and variance, whereas Pearson correlation measures the strength of linear association between inferred and true values and is insensitive to scaling and offset. Reporting both is useful because both fitness values and ranking are used in evolutionary studies [2, 10].

#### Z-score calibration

We computed a standardized error for each parameter,

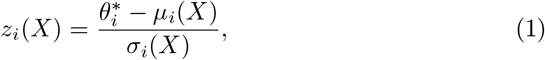

where *σ*_*i*_(*X*) is the posterior standard deviation. Under a well-calibrated posterior, these Z-scores should approximately follow a standard normal distribution, *z*_*i*_ ∼ *𝒩* (0, 1). Accordingly, a mean near zero indicates little systematic bias, a standard deviation near one indicates appropriate uncertainty quantification, and a mean absolute value near 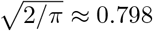 is consistent with overall calibration. Parameters with| mean(*z*) | > (1.0) were flagged as potentially biased, and those with std(*z*) > 1.5 were flagged as potentially overconfident.

#### Posterior contraction

To assess whether NPE updates its beliefs beyond the prior 𝒩 (0, 0.2), we compared the prior with the learned posterior KDE aggregated across all held-out test cases for each chromosome-arm parameter. Consistent narrowing of the posterior relative to the prior indicates that the model extracts information from the clonal CNA observations rather than merely reproducing prior uncertainty.

## 3 Results and Discussion

We evaluated the learned posterior on held-out simulated tumors using the diagnostics described in Section 2.6.

### Posterior calibration of CloneMLP-NPE

Figures 1 and 2 show the posterior Z-score distributions for all 44 chromosome-arm parameters under **CloneMLP-NPE**. Across most arms, the Z-score distributions are approximately symmetric and centered near zero, indicating little systematic bias in the posterior mean estimates. The distributions are generally contained within the reference interval [− 2, 2], with only moderate tails for a small number of parameters. Mean absolute Z-scores ranged from approximately 0.60 to 0.80 (Figure 3), with most values lying below the theoretical expectation of 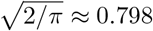 under 𝒩 (0, 1). Overall, these results indicate that the posterior learned by **CloneMLP-NPE** is well calibrated, with at most mild underconfidence for several chromosome arms. No arm showed strong evidence of systematic bias or severe overconfidence under our diagnostic thresholds.

**Fig. 1.**
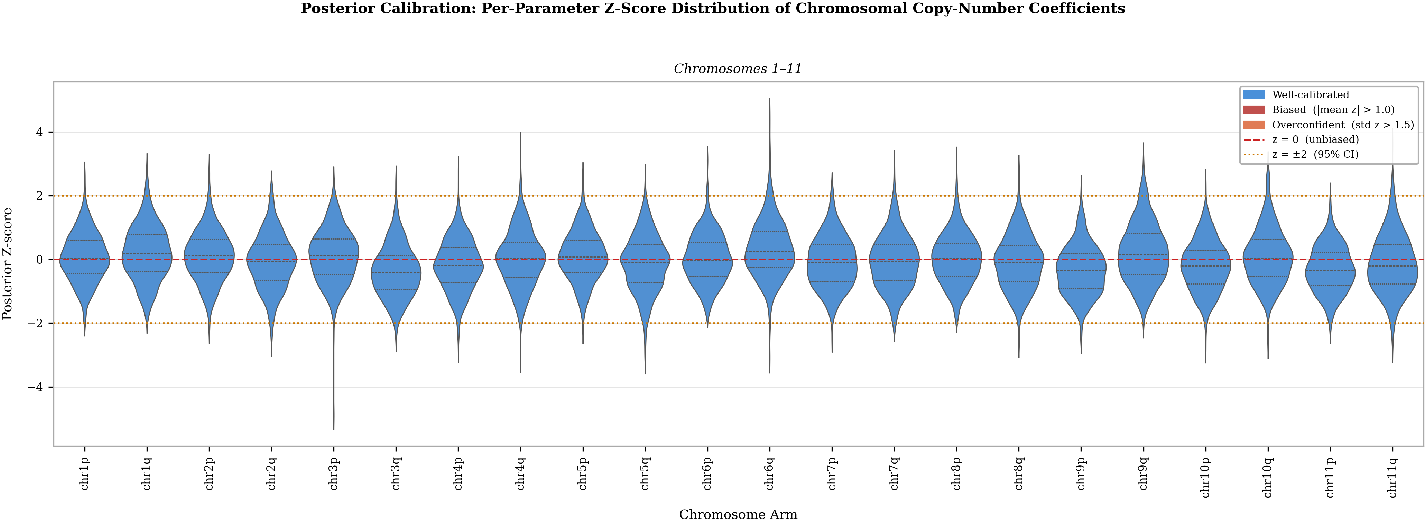
Posterior Z-score distributions for chromosome arms 1p–11q across held-out simulations under **CloneMLP-NPE**. Violin widths represent density, and internal lines indicate the interquartile range and median. The red dashed line marks *z* = 0, and the orange dotted lines mark *z* = ±2. Most distributions are approximately centered near zero, consistent with good posterior calibration.

**Fig. 2.**
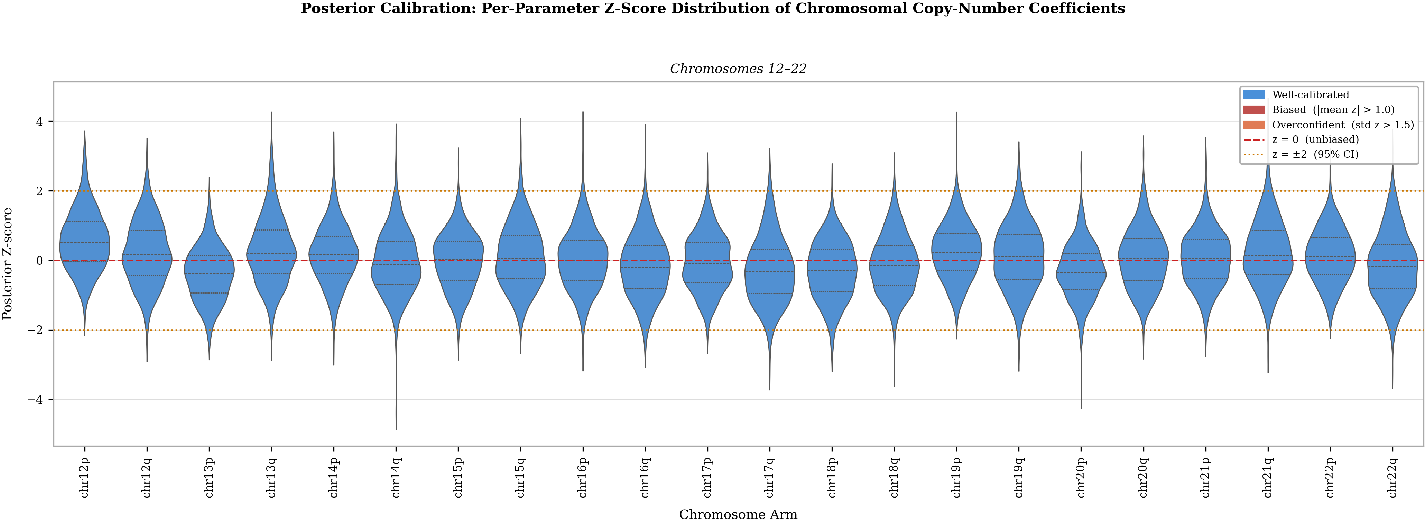
Posterior Z-score distributions for chromosome arms 12p– 22q across held-out simulations under **CloneMLP-NPE**. Layout and reference lines are the same as in Figure 1. Most distributions remain approximately centered near zero, indicating broadly consistent calibration across parameters.

**Fig. 3.**
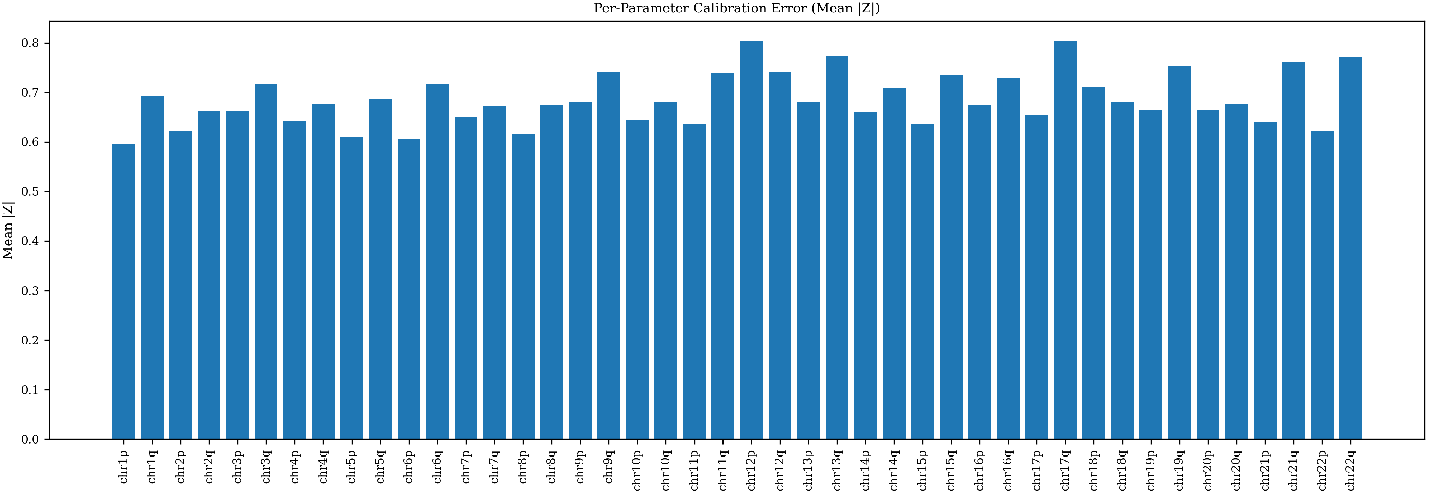
Mean absolute Z-score for each chromosome-arm parameter across held-out simulations under **CloneMLP-NPE**. Most values lie close to the theoretical expectation 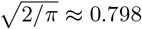 under a standard normal distribution, indicating generally good posterior calibration with mild underconfidence for several parameters.

### Posterior mean recovery of CloneMLP-NPE

Figures 4 and 5 compare the true chromosome-arm selection coefficients 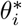with posterior means *µ*_*i*_(*X*) across held-out simulations for **CloneMLP-NPE**. The posterior means exhibit a clear positive relationship with the ground-truth values for all arms, and the fitted regression lines broadly follow the ideal *y* = *x* diagonal, although with shallower slopes that indicate some shrinkage toward zero. Recovery quality varies across chromosome arms, with *R*^2^ values ranging from 0.34 to 0.62. The strongest recovery is observed for arms such as chr2p and chr13p (*R*^2^ ≈ 0.62), whereas weaker but still substantial recovery is seen for arms such as chr17q and chr22q (*R*^2^ ≈ 0.36 and 0.34, respectively). These plots suggest that **CloneMLP-NPE** is able to recover a substantial fraction of the variation in the true parameters, although posterior means remain somewhat compressed toward the prior mean, especially for larger-magnitude coefficients.

**Fig. 4.**
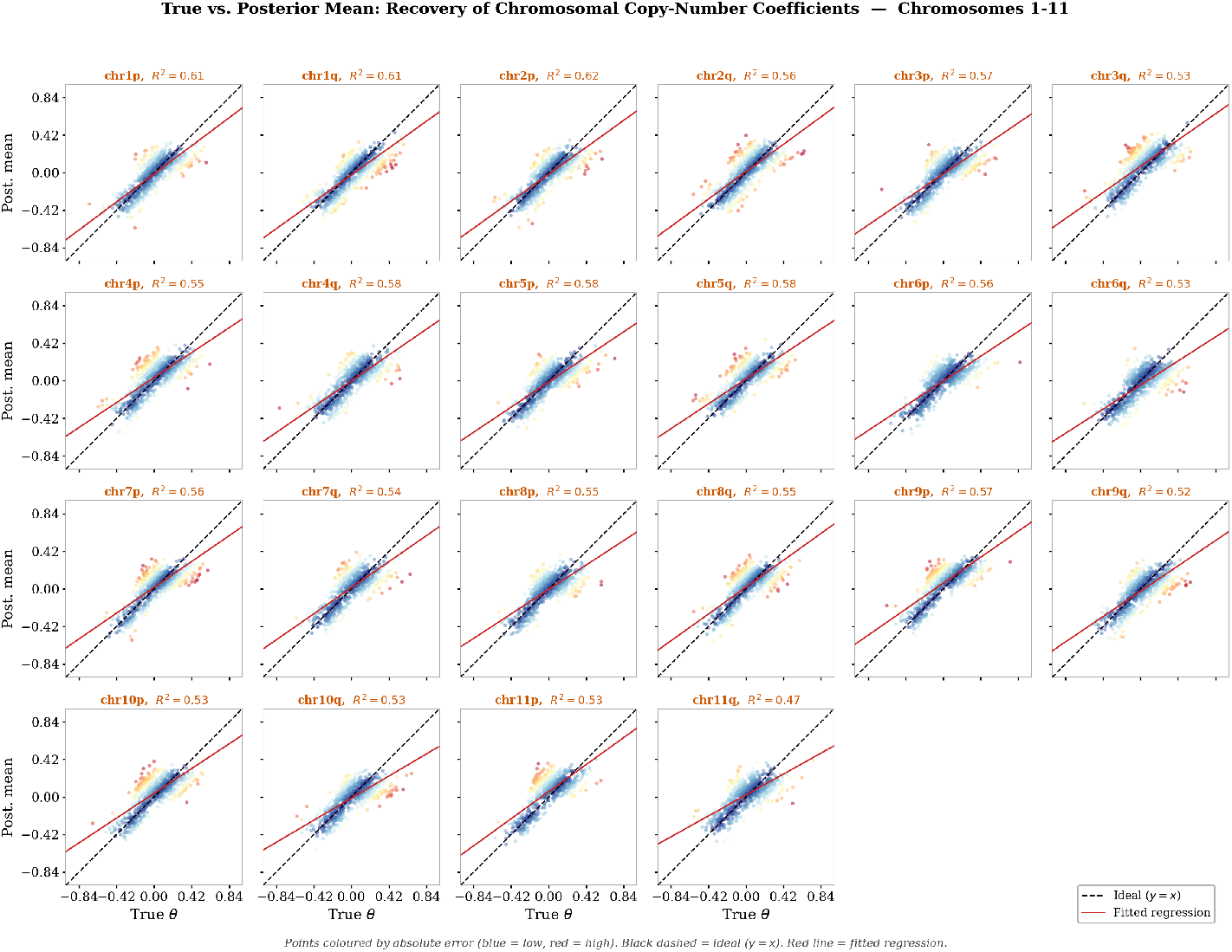
True selection coefficients versus posterior means for chromosome arms 1p–11q across held-out simulations under **CloneMLP-NPE**. Points are colored by absolute error (blue: low, red: high). The black dashed line denotes the ideal relation *y* = *x*, and the red solid line shows the fitted regression. Annotated *R*^2^ values summarize posterior mean recovery for each arm.

**Fig. 5.**
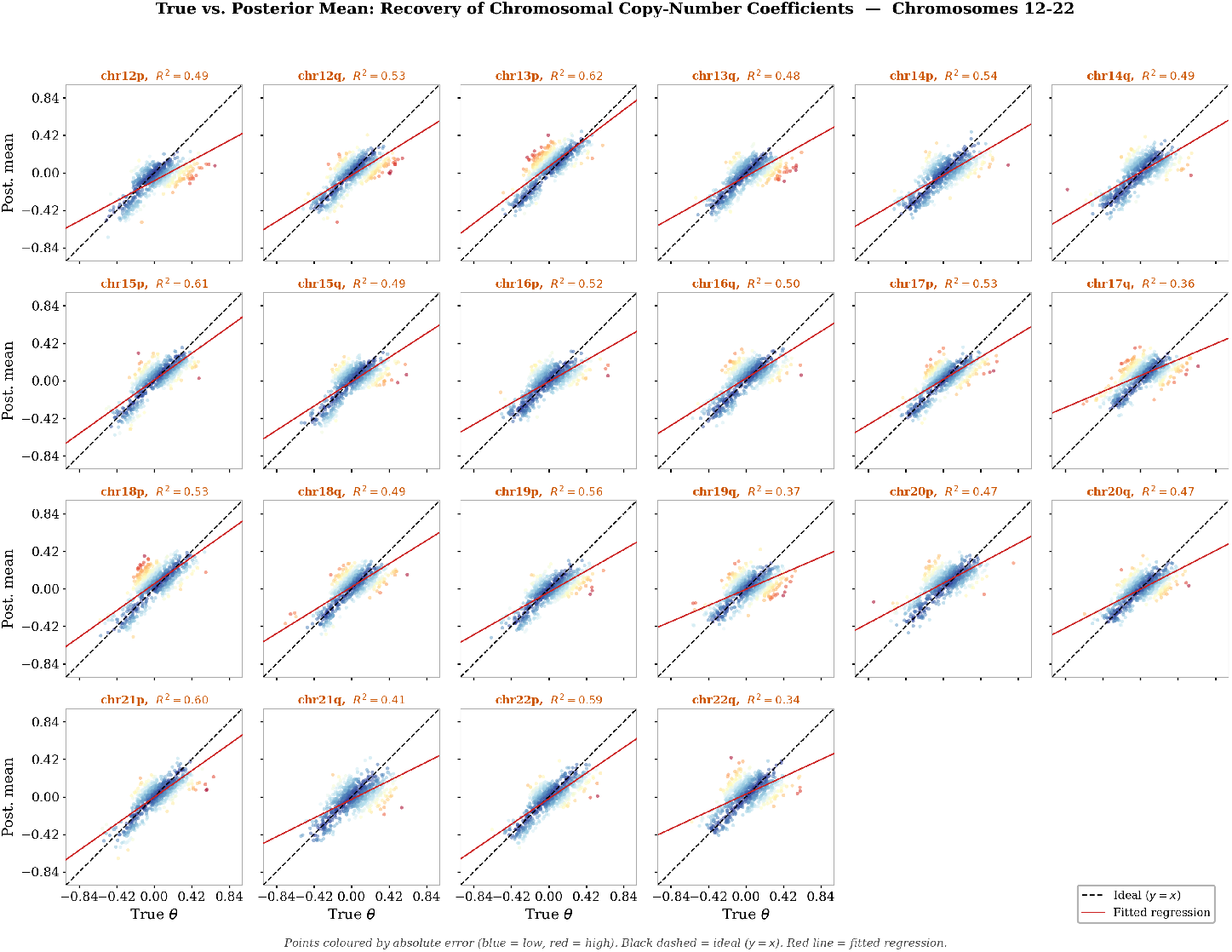
True selection coefficients versus posterior means for chromosome arms 12p– 22q across held-out simulations under **CloneMLP-NPE**. Layout and annotations are the same as in Figure 4. Regression lines generally follow the ideal diagonal, indicating moderate to strong recovery across most parameters.

### Comparison across models

Table 1 summarizes posterior mean recovery for the six chromosome arms with the highest *R*^2^ in **CloneMLP-NPE. CloneMLP-NPE** consistently outperformed both baselines on all six arms, achieving Pearson correlations of approximately 0.77–0.79 and corresponding *R*^2^ values near 0.60. **DominantClone-NPE** showed intermediate performance, while **CloneAtt-NPE** was weakest overall. This pattern suggests that using the full multi-clone CNA representation is more informative than relying only on the largest clone, and that in this setting the simpler MLP encoder is more effective than the Set Transformer for extracting features relevant to posterior inference.

**Table 1.** Comparison of posterior mean recovery across the three models for the six chromosome arms with the highest *R*^2^ in CloneMLP-NPE model. For each model, *R*^2^ is computed as the square of the Pearson correlation coefficient.

| Chromosome arm | CloneMLP-NPE |  | CloneAtt-NPE |  | DominantClone-NPE |  |
| --- | --- | --- | --- | --- | --- | --- |
| | $R^2$ | Pearson $r$ | $R^2$ | Pearson $r$ | $R^2$ | Pearson $r$ |
| chr13p | 0.617 | 0.786 | 0.042 | 0.206 | 0.098 | 0.313 |
| chr2p | 0.616 | 0.785 | 0.161 | 0.401 | 0.346 | 0.589 |
| chr1q | 0.614 | 0.784 | 0.015 | 0.122 | 0.253 | 0.503 |
| chr1p | 0.608 | 0.780 | 0.036 | 0.190 | 0.322 | 0.567 |
| chr15p | 0.606 | 0.778 | 0.024 | 0.155 | 0.119 | 0.345 |
| chr21p | 0.600 | 0.772 | 0.008 | 0.091 | 0.145 | 0.381 |

## 4 Conclusion

We presented a likelihood-free Bayesian framework for inferring chromosome-arm selection coefficients from clonal CNA profiles using neural posterior estimation with a normalizing-flow posterior and tumor representations derived from clonal CNA summaries. Among the three models considered, **CloneMLP-NPE** produced the strongest overall results. Using the whole-tumor CNA matrix, **CloneMLP-NPE** yielded well-calibrated posteriors across all 44 chromosome arms, with Z-score distributions that were approximately symmetric and centered near zero, indicating little systematic bias. Posterior mean recovery was consistently positive, with moderate-to-strong performance across chromosome arms, showing that the model extracts substantial signal from clonal CNA observations. Comparison with **CloneAtt-NPE** and **DominantClone-NPE** further showed that **CloneMLP-NPE** achieved the strongest posterior mean recovery on the top-performing parameters.

These results suggest that whole-tumor CNA representations are informative for selection inference, while also indicating that some effects remain only partially identifiable under the current simulation and observation setting. Future work will expand the simulation dataset and broaden prior support to better cover larger-magnitude selection coefficients. In addition, although **CloneAtt-NPE** underperformed in the current study, its permutation-invariant architecture remains a promising approach for representing heterogeneous clonal populations. Further studies involving training **CloneAtt-NPE** on larger datasets and further refinement of its architecture are required to assess whether improved set-based representations can yield stronger inference performance.

## Acknowledgments

This study was partially supported by NSF grants 2415564 and 2415562. The author have no competing interests to declare.

## Notes

### Competing Interest Statement

The authors have declared no competing interest.

## References

1. Cranmer, K., Brehmer, J., Louppe, G.: The frontier of simulation-based inference. Proceedings of the National Academy of Sciences 117(48), 30055–30062 (2020)

2. Crona, K., Gavryushkin, A., Greene, D., Beerenwinkel, N.: Inferring genetic interactions from comparative fitness data. Elife 6, e28629 (2017)

3. Lee, J., Lee, Y., Kim, J., Kosiorek, A., Choi, S., Teh, Y.W.: Set transformer: A framework for attention-based permutation-invariant neural networks. In: International conference on machine learning. pp. 3744–3753. PMLR (2019)

4. Luo, X.G., Kuipers, J., Rupp, K., Takahashi, K., Beerenwinkel, N.: Bayesian inference of fitness landscapes via tree-structured branching processes. Bioinformatics 41(Supplement_1), i160–i169 (2025)

5. Lynch, A.R., Arp, N.L., Zhou, A.S., Weaver, B.A., Burkard, M.E.: Quantifying chromosomal instability from intratumoral karyotype diversity using agent-based modeling and bayesian inference. Elife 11, e69799 (2022)

6. Nowell, P.C.: The clonal evolution of tumor cell populations: Acquired genetic lability permits stepwise selection of variant sublines and underlies tumor progression. Science 194(4260), 23–28 (1976)

7. Papamakarios, G., Nalisnick, E., Rezende, D.J., Mohamed, S., Lakshminarayanan, B.: Normalizing flows for probabilistic modeling and inference. Journal of Machine Learning Research 22(57), 1–64 (2021)

8. Papamakarios, G., Sterratt, D., Murray, I.: Sequential neural likelihood: Fast likelihood-free inference with autoregressive flows. In: The 22nd international conference on artificial intelligence and statistics. pp. 837–848. PMLR (2019)

9. Salehi, S., Kabeer, F., Ceglia, N., Andronescu, M., Williams, M.J., Campbell, K.R., Masud, T., Wang, B., Biele, J., Brimhall, J., et al.: Clonal fitness inferred from time-series modelling of single-cell cancer genomes. Nature 595(7868), 585–590 (2021)

10. Skums, P., Tsyvina, V., Zelikovsky, A.: Inference of clonal selection in cancer populations using single-cell sequencing data. Bioinformatics 35(14), i398–i407 (2019)

11. Stadler, T., Pybus, O.G., Stumpf, M.P.: Phylodynamics for cell biologists. Science 371(6526), eaah6266 (2021)

12. Tejero-Cantero, A., Boelts, J., Deistler, M., Lueckmann, J.M., Durkan, C., Gonçalves, P.J., Greenberg, D.S., Macke, J.H.: Sbi–a toolkit for simulation-based inference. arXiv preprint arXiv:2007.09114 (2020)

13. Tijhuis, A.E., Foijer, F.: Characterizing chromosomal instability-driven cancer evolution and cell fitness at a glance. Journal of Cell Science 137(1), jcs260199 (2024)

14. Weiner, S., Bansal, M.S.: Sistem: simulation of tumor evolution, metastasis, and dna-seq data under genotype-driven selection. Bioinformatics 41(12), btaf634 (2025)

15. Williams, M.J., Werner, B., Heide, T., Curtis, C., Barnes, C.P., Sottoriva, A., Graham, T.A.: Quantification of subclonal selection in cancer from bulk sequencing data. Nature genetics 50(6), 895–903 (2018)

16. Xiang, Z., Liu, Z., Dinh, K.N.: Inference of chromosome selection parameters and missegregation rate in cancer from dna-sequencing data. Scientific Reports 14(1), 17699 (2024)

